## Supplemental Figures 1-6 for "Lots of movement, little progress: A review of reptile home range literature" (from another version of this preprint: version 1, 2020-12-04)

**Supplementary Materials**

Includes Table S1 Figures S1-S6

***Table S1***. List of other methods recorded.

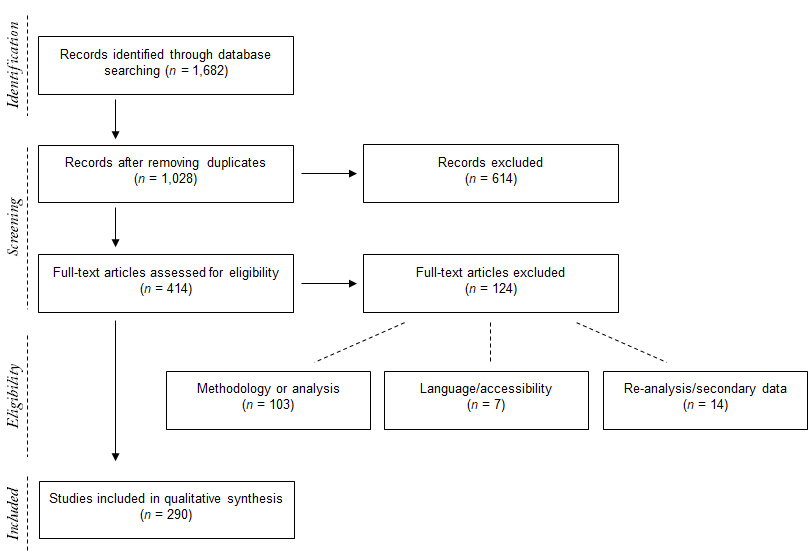

***Figure S1.*** *Preferred Reporting Items for Systematic Reviews and Meta-Analyses* *(PRISMA) flowchart (Moher et al. 2009) for systematic review of articles on reptile biotelemetry studies*.

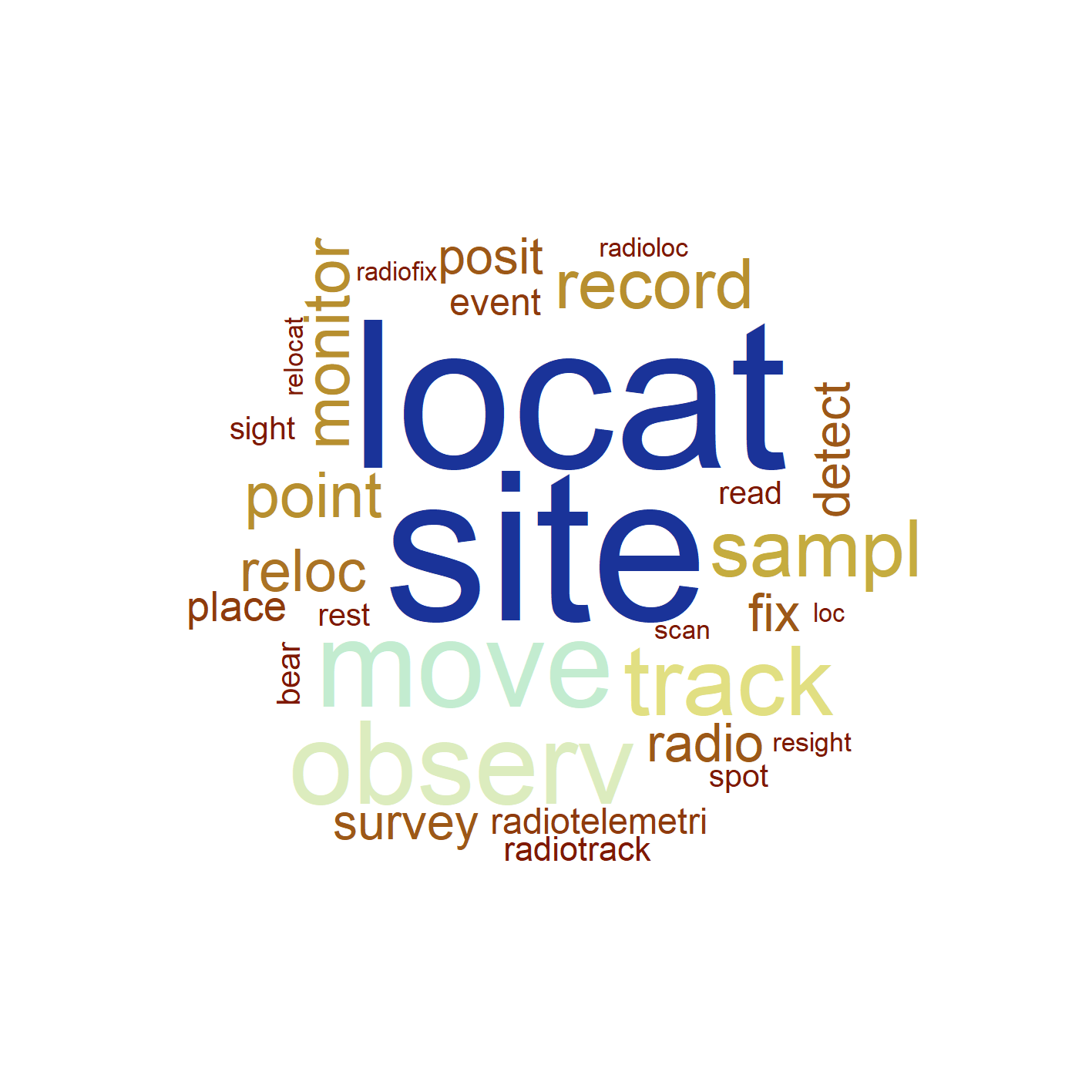

***Figure S2.*** *Word cloud showing the stem of terms used at least twice to describe tracking an animal, colour and size denote frequency of use.*

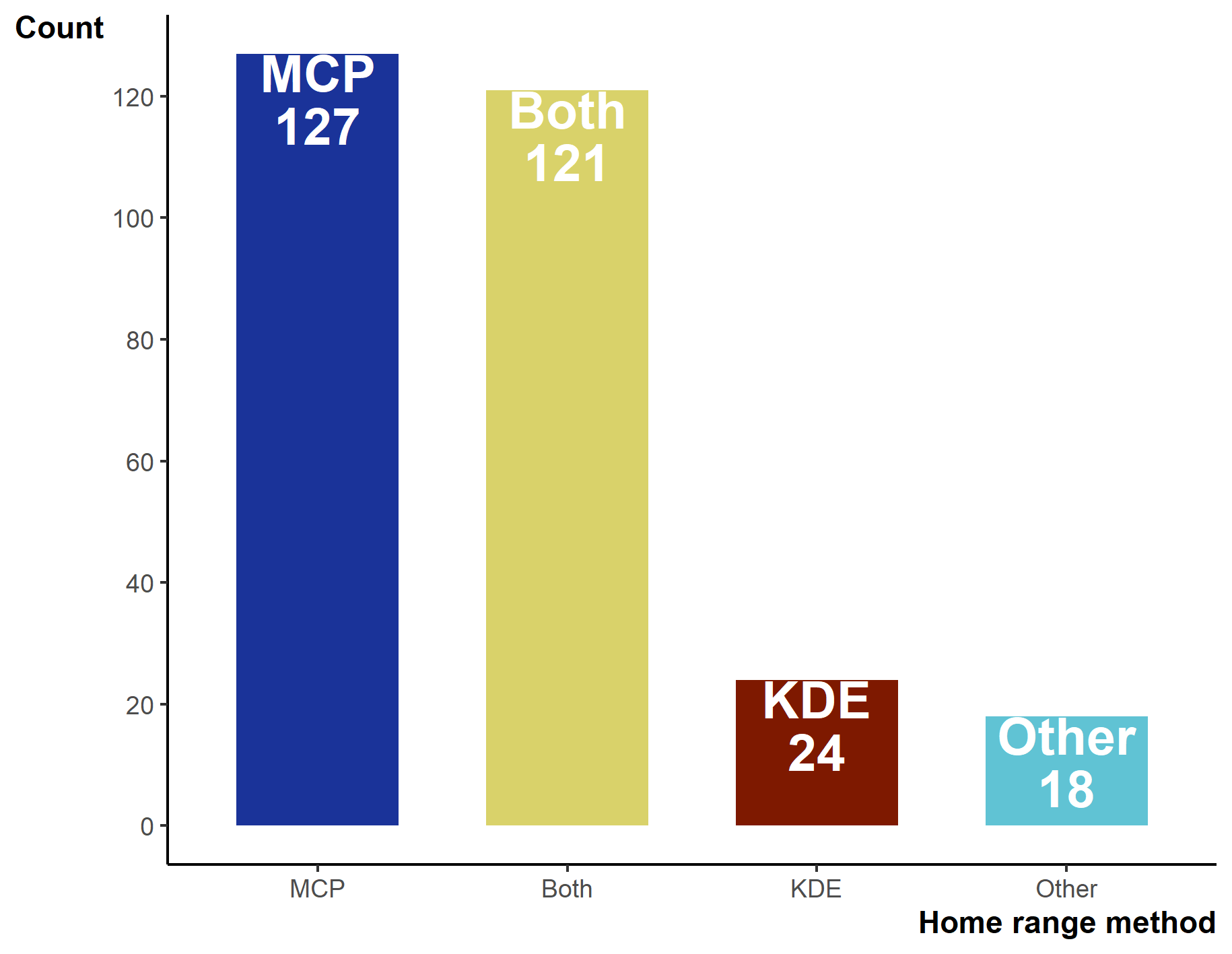

***Figure S3.*** *The total number of studies using Minimum Convex Polygon (MCP), Kernel Density Estimations (KDE), or both. “Other” only includes studies that did not use either MCPs or KDEs.*

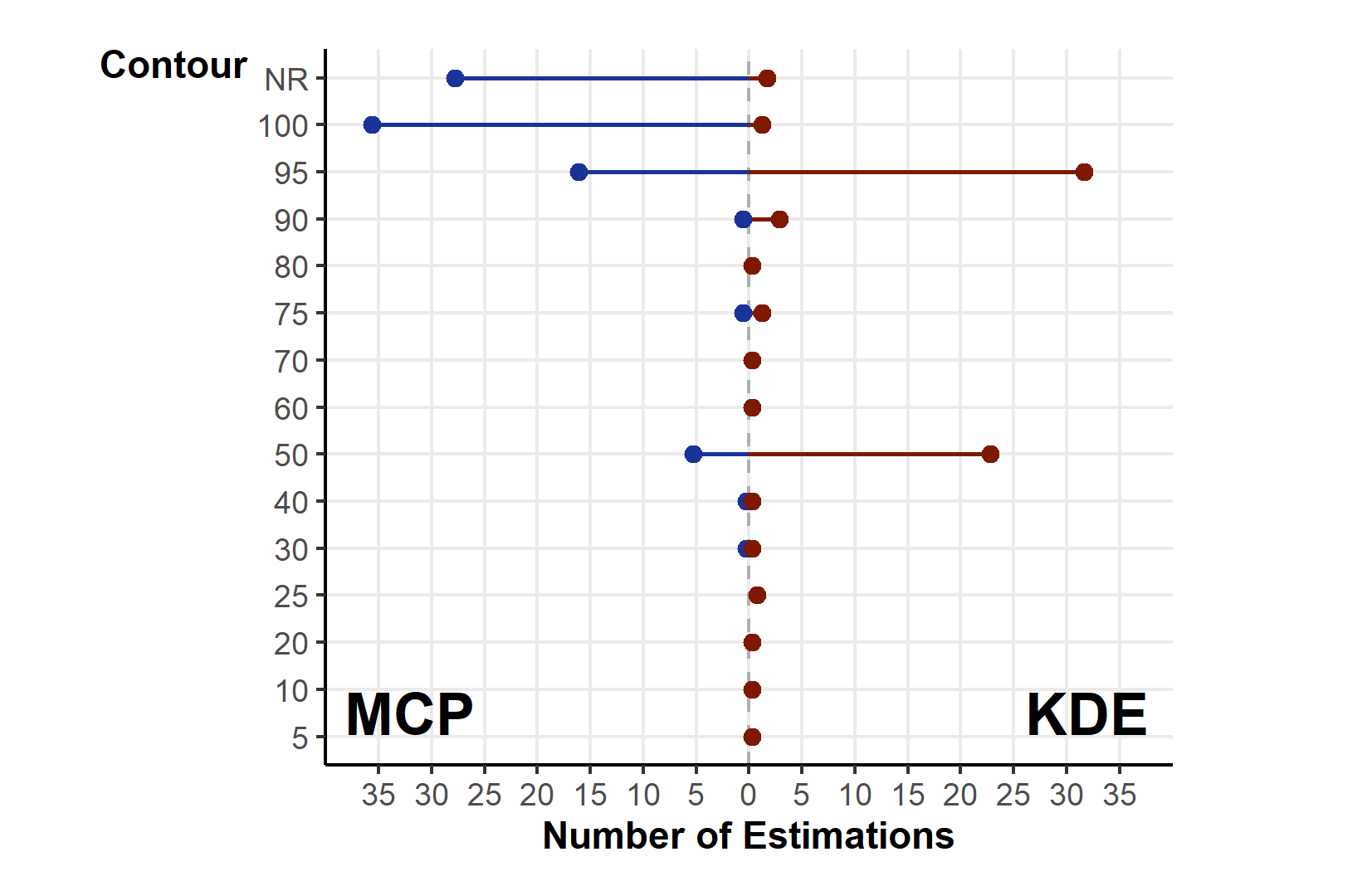

***Figure S4.*** *The number of times a given contour was used in conjunction with Minimum Convex Polygon (MCP), and Kernel Density Estimations (KDE). Unused contours have been excluded from the y-axis*.

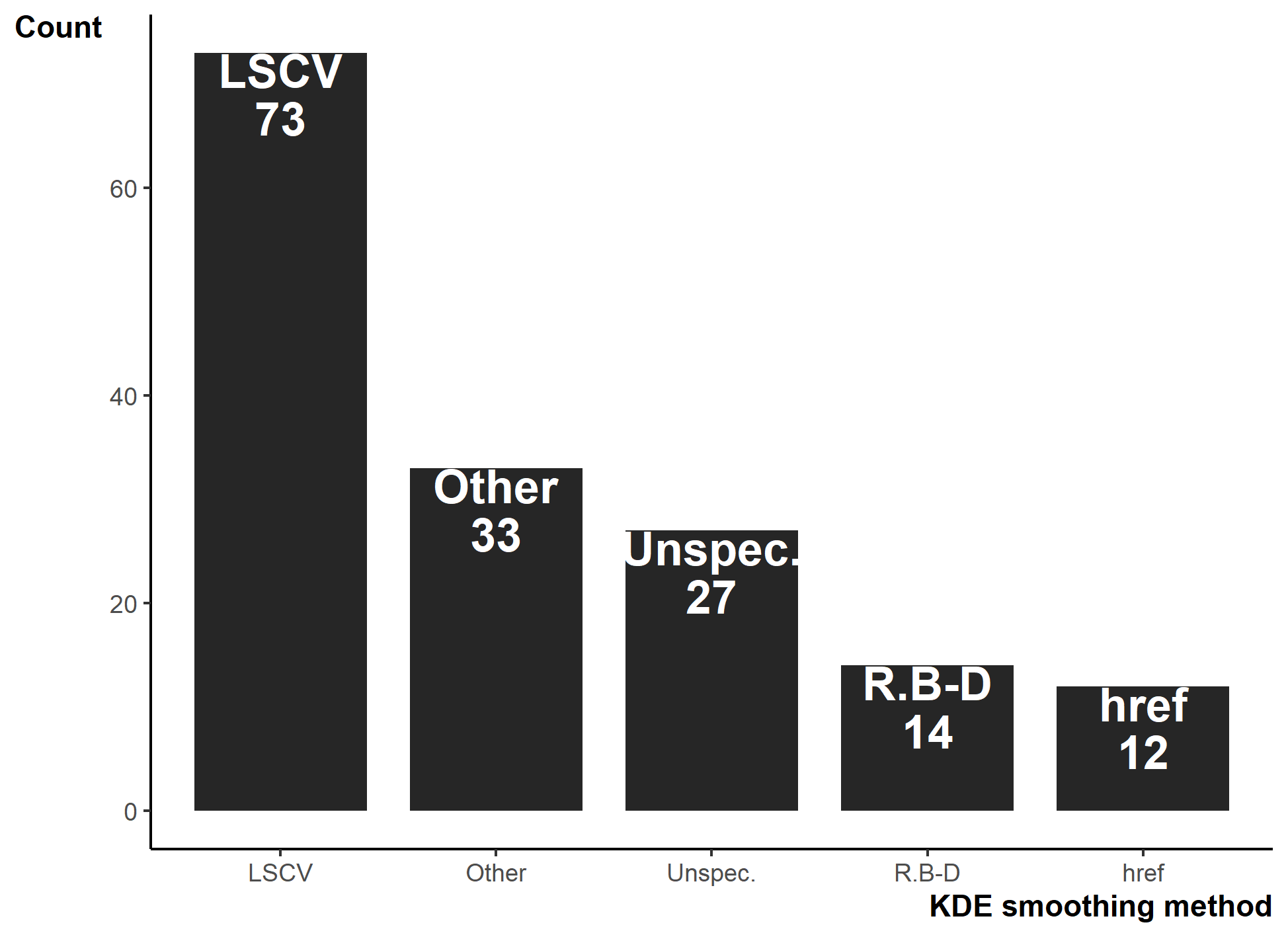

***Figure S5****. Smoothing factors used for KDE. Unspec. are instances where the smoothing factor (h) was not explicitly stated. R.B-D are cases where studies manipulated h to ensure KDE area output equalled MCP area estimation as suggested by Row and Blouin-Demers 2006.*

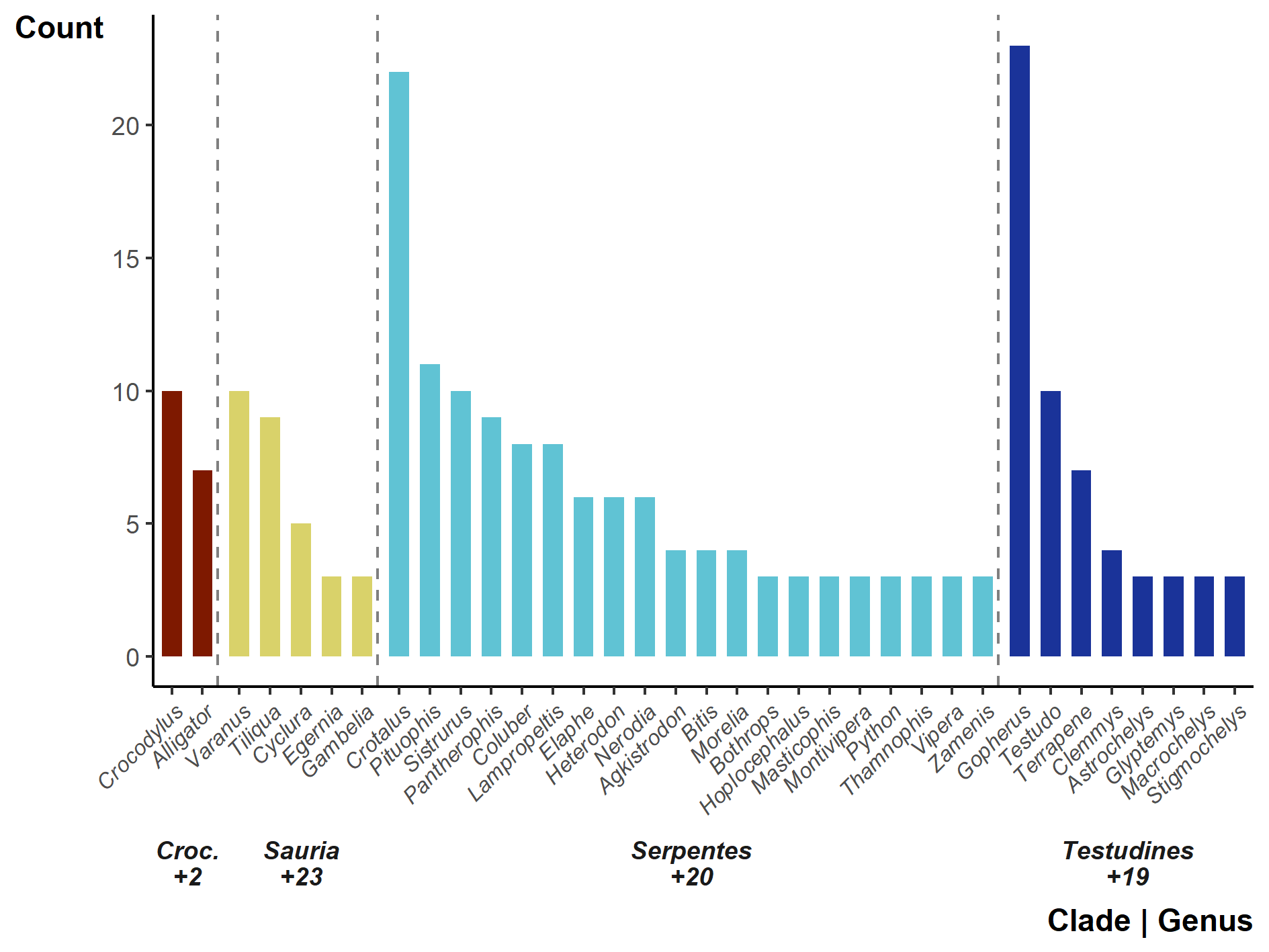

***Figure S6.*** *The number of studies by genus and clade, provided the genus appears in three or more studies. Numbers below clades indicate the number of genera not listed (i.e., genus only appearing in two or fewer studies).*
